# IMPORTANT: Advanced Pollen Classification of Indian Medicinal Plants through SEM and Computer Vision

**DOI:** 10.1101/2025.01.08.631879

**Authors:** Jaidev Sanjay Khalane, Nilesh D. Gawande, Shanmuganathan Raman, Subramanian Sankaranarayanan

## Abstract

Pollen grains of plant species have unique morphological characteristics. The variability in shape, size, and microscopic pollen surface features can be efficiently used to determine the plant species to which they belong. This approach can be instrumental in regions with rich biodiversity in plant species, specifically in medicinal plant. The creation of a pollen dataset for these species using SEM images and a computer vision application can be beneficial for their identification. We have developed a robust approach utilizing scanning electron microscopy (SEM) to generate high-resolution pollen images from 28 plant species of medicinal importance and employed computer vision techniques for accurate segmentation and classification based on diverse morphological features. In this study, a dataset comprising 269 images for segmentation and 5842 images was used for classification across 28 classes. In addition, we have created a globally accessible database named IMPORTANT (Indian Medicinal Plants Pollen Images Dataset for Research, Training, and Analysis of Neural Networks) to facilitate easy retrieval and sharing of pollen images. This study can effectively identify medicinal plant species by utilizing the microscopic features of pollen SEM images through the IMPORTANT database. Since the pollen of even closely related species vary in their morphological characteristics, this approach can efficiently determine the closely related species with accuracy and precision.

## Introduction

Pollen grains are microscopic entities in the angiosperms, containing microgametophytes essential for the production of male gametes in plant reproduction [1]. The diverse pollen morphology is a common phenomenon in angiosperms, which is helpful in the identification and classification of species. Pollen has a wide range of applications, including taxonomic classification, examining pollen allergies, historical climate change analysis, and archaeological findings [2,3].

Pollen have unique morphology that differs from species to species and includes shapes such as boats, cups, prisms, spheroids, and many more. The overall structure of pollen results from pollen shapes and the arrangement of surface furrows that facilitate pollination [4]. In addition, the outer layer of the pollen wall, known as the exine, and surface features, along with surface furrows, contribute to the unique complex structures of pollen, facilitating their taxonomic classification [5,6]. Various microscopy techniques, such as light microscopy, scanning electron microscopy (SEM), and transmission electron microscopy TEM) can capture the structural feature details of the pollen. However, SEM facilitates a more accurate determination of pollen surface morphology [7]. The SEM images captured for the closest species of *Bougainvillea*, such as *B. alba, B. spectabilis*, and *B. buttiana* can efficiently differentiate these species based on the pollen surface morphology characteristics [8].

Various databases have been created for the taxonomic identification of plant species by pollen images captured using microscopy techniques with varying rates of accuracy and precision. For example, the dataset POLLEN73S, consisting of 2523 images captured using light microscopy from 73 pollen types in the Brazilian Savanna exhibited higher precision and recall of 95.7% [9]. In this study, pollen were classified using CNN-based models such as DenseNet-201 and ResNet-50. Another study included the dataset from the light microscopy captured images of pollen across 40 species. This study extensively used a deep learning (DL) approach to automate and enhance pollen grain classification, especially in the Great Basin Desert, Nevada, USA. The transfer learning on 10,000 images with pre-trained Convolutional Neural Networks (CNNs), including AlexNet, VGG-16, MobileNet-V2, ResNet, ResNeSt, SE-ResNeXt, and Vision Transformer (ViT) demonstrated that ResNeSt-110 model outperformed other CNNs in various parameters including accuracy (97.24%), precision (97.89%), F1 score (96.86%) and recall (97.13%) [10]. These studies demonstrate that DL models with transfer learning can significantly improve the accuracy and speed of pollen classification. An annotated dataset of types of Brazilian Savannah pollen consisted of 805 pollen images captured using light microscopy of 23 species [11]. The study establishes a baseline for human and computer performance, implementing and fine-tuning three engineered feature extractors and four machine learning techniques. Another dataset, Cretan Pollen Dataset v1 (CPD-1), had 4,034 segmented pollen grains from 20 species collected in Crete, Greece. In this study, 400x magnification images were captured using light microscopy [12].

Despite the availability of robust imaging techniques, there is still a lack of extensive large datasets specialized for computer vision applications, including segmentation and classification, particularly in Indian Medicinal Plants. Given this context, we have developed a comprehensive scanning electron microscope (SEM)-based dataset to support segmentation and detection in computer vision applications across 28 Indian medicinal plant species. In addition, we analyzed the performance of the YOLOv11 segmentation model [13] for the separation of pollen from background noise in images and tested various classification models based on Convolutional Neural Networks, including ResNet50 [14], VGG16 [15], MobileNet [16], AlexNet [17], and the Vision Transformers [18] to classify segmented pollen images by species. Furthermore, we explored similarities and differences in the pollen images across species within the same families. This study provides a comprehensive SEM pollen images dataset for Indian medicinal plant species and allows the identification of plants with accuracy and precision.

## Material and methods

### Dataset collection and characterization of pollen surface morphology

The dataset included the pollen SEM images of 28 distinct flower species with medicinal properties collected from the Indian Institute of Technology Gandhinagar (IITGN) campus in Gujarat, India. Pollen surface morphology for the species was determined using a field emission scanning microscope (FE-SEM, JEOL^®^ JSM-7600F) at the Central instrumentation facility at IITGN. The database named IMPORTANT (Indian Medicinal Plants Pollen Image Database for Research, Training, Analysis, and Neural Networks Testing), which contains details of the species and pollen SEM images, was created. The details of plant species used for capturing the pollen SEM images of pollen and their families are provided in **Supplementary Table 1**.

### Computational methodology

The collected dataset was used for the training and testing of various computer vision models specialized in segmentation, facilitating the identification and segmentation of pollen from the background images and classifying them into respective species.

### Segmentation model

The segmentation model YOLOv11 enables instance segmentation by accurately identifying and segmenting individual objects within images. To segment the pollen from the image (corresponding to the class “pollen”), the lighter version of YOLOv11, YOLOv11n, was used. The model was trained for 20 epochs with a batch size of 8. The AdamW optimizer was used with a learning rate of 0.002 and a momentum of 0.9. Out of 269 images, 32 images were further considered for validation. The number of classes present was one, corresponding to the class “pollen.” The distribution of the segmentation dataset across species is detailed in **Supplementary Table 2**. The performance metrics used for the evaluation of the segmentation model included mask precision (Number of correctly segmented pixels/Total number of segmented pixels), mask recall (Number of correctly segmented pixels/Total number of true instance pixels), box precision (Intersection over the Union of the predicted bounding box and the ground truth bounding box/Area of the predicted bounding box), and box recall (Intersection over the Union of the predicted bounding box and the ground truth bounding box/Area of the true bounding box).

### Classification model

The classification task involves the classification of the given images of the pollen into their respective species. This study used four types of CNN-based models and a transformer-based model. The input for the model consisted of images resized to 224. For the CNN based models, such as ResNet 50, VGG 16, MobileNet and AlexNet, the Adam optimiser with a learning rate of 0.001 and a decay of 1e^-6^ was employed. In the transformer based model, Adam optimiser with a learning rate was 1e^-4^ was used. The categorial cross entropy and accuracy were used as the objective functions and loss metric under consideration, respectively. In this study, approximately 80% of the entire dataset was used for training, while the rest was used equally for testing and validation. The training and validation batch size consisted of 100 images, and the entire training was run for 25 epochs. The training process was carried out using hardware Nvidia Quadro RTX 5000 (16GB). In the classification, the segmented images were augmented with 2D affine transformations. The classification dataset consists of a total of 5962 images distributed across 28 classes (species). The details for the distribution of classification of the dataset across species are given in **Supplementary Table 3**. The confusion matrices for the test dataset consisting of 489 images were generated, and the classification report, which included classwise precision, recall, and F1 scores, was determined. Comparison between the performances of the models in terms of accuracy, macro average precision, macro average recall, and macro average F1 scores was assessed. Other models used included VGG 16, MobileNet, AlexNet, and ViT. The performance metrics used for the evaluation of the classification model included accuracy (True Positives + True Negatives/Total Instances), precision (True Positives/(True Positives + False Positives), recall (True Positives/(True Positives + False Negatives), and F1 score (2×Precision×Recall/(Precision + Recall).

### Python libraries used in the study

The models were trained and implemented using Python (https://www.python.org/). Other libraries such as TensorFlow [19], Keras [20], Transformers [21], PyTorch [22], with NumPy [23], Pandas [24], Seaborn [25], Matplotlib [26], Scikit Learn [27] and TorchVision [28] were used. The library OpenCV [29] was used for data preprocessing and visualization.

## Result and Discussion

In this study, we have developed a database of pollen SEM images from 28 medicinal plant species for their precise and accurate identification, as illustrated in **Fig. 1**.

**Fig. 1.**
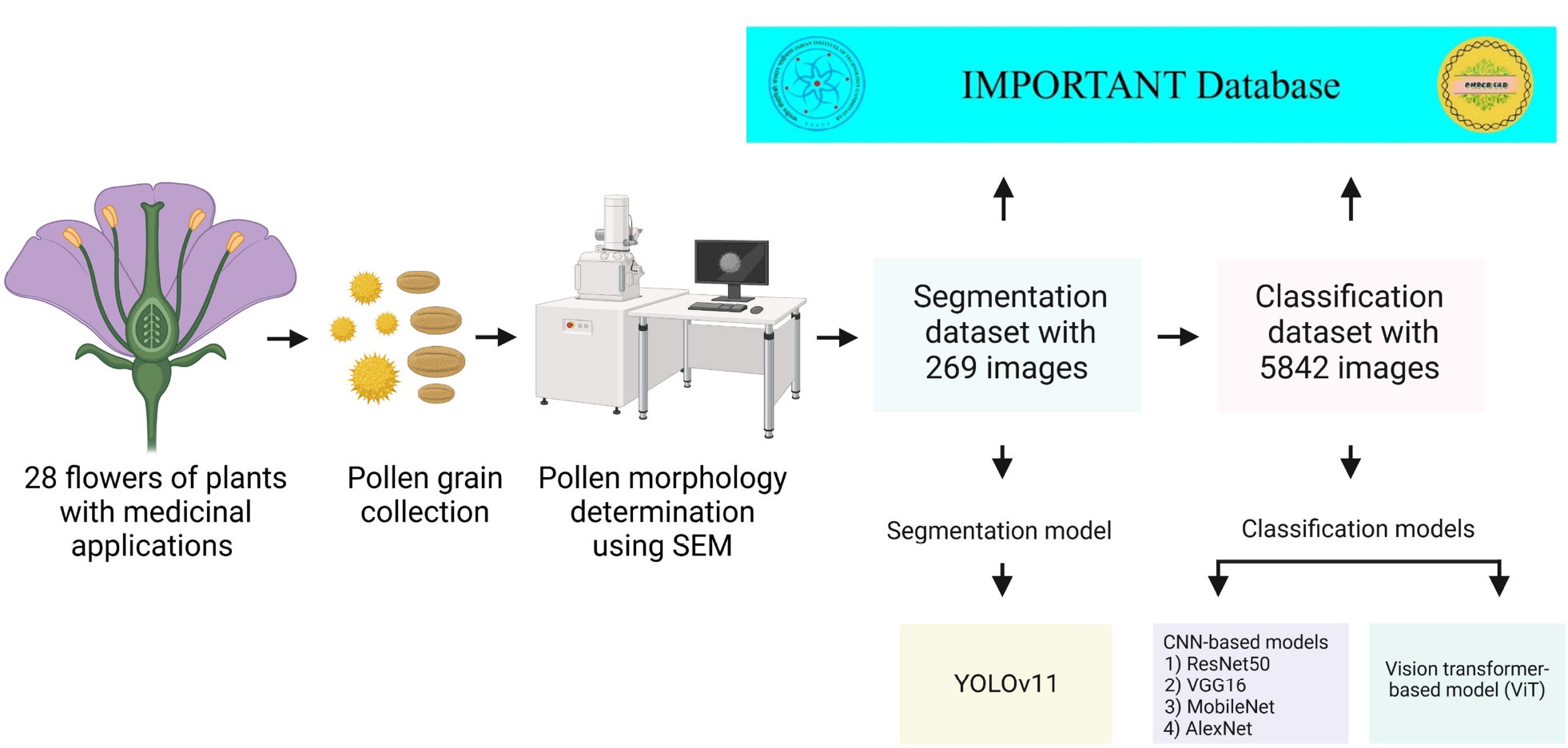
Characterization of pollen surface morphology and construction of datasets for pollen SEM images. The pollen surface morphology for 28 medicinal plant species was determined using a field emission scanning microscope (FE-SEM). The YOLOv11 segmentation model was used to separate pollen from background noise. Various CNN-based classification models (ResNet50, VGG16, MobileNet, and AlexNet) and Vision Transformers (ViT) were used for the classification of pollen into respective species. The species uses 269 and 5842 datasets for the segmentation and classification datasets, respectively.

### Pollen for the species exhibit distinct and unique morphological characters

The pollen images captured using SEM showed distinct and unique morphological characteristics in terms of the shape and the pollen surface. The pollen shapes for the species varied from spheroidal to irregular shapes. The pollen surface morphology for almost all the species exhibits distinct characteristics, including smooth textures, spiny surfaces, and surfaces with dots. The pollen SEM images are given in **Fig. 2**, and the details for the shapes and surface morphology of the pollen are provided in **Table 1**. The high-resolution images obtained through SEM provide extensive details of the pollen surface architecture, which are crucial for accurate species identification and classification. Our study confirms the effectiveness of SEM in capturing detailed pollen grain morphology, which is essential for distinguishing between closely related species [8]. The precise delineation of exine patterns, apertures, and surface ornamentations facilitates precise morphological comparisons.

**Table 1.** Details of the pollen shapes and surface morphology for plant species.

| Plant species | Pollen shape | Surface texture |
| --- | --- | --- |
| *Azadirachta indica* | Cuboidal sphere | Slightly rough |
| *Caesalpinia pulcherrima* | Spherical | Smooth |
| *Canna indica* | Spheroidal | Slightly rough |
| *Cassia fistula* | Prolate spheroidal | Dotted |
| *Cassia javanica* | Convex triangular | Smooth |
| *Catharanthus pusillus* | Cuboidal sphere | Smooth |
| *Catharanthus roseus* (Linn.) Don | Cubical sphere | Slightly rough |
| *Cordia sebestena* | Prolate spheroidal | Densely reticulated |
| *Delonix regia* | Prolate spheroidal | Reticulated with large dots |
| *Eclipta prostrata* | Spheroidal | Spiny |
| *Gaillardia pulchella* | Spheroidal | Spiny with dotted holes |
| *Glandularia bipinnatifida* | Irregular | Regular |
| *Helianthus annuus* | Spheroidal | Spiny |
| *Hibiscus arnottianus* | Spheroidal | Slightly spiny |
| *Hibiscus rosa-sinensis* | Spheroidal | Spiny |
| *Hymenocallis littoralis* | Cubical sphere | Loosely reticulated |
| *Ixora coccinea* | Irregular | Rough |
| *Jasminum sambac* | Irregular | Rough |
| *Murraya paniculata* | Oblate spheroidal | Densely reticulated |
| *Ocimum tenuiflorum* | Spheroidal | Loosely reticulated with large dots |
| *Ocimum gratissimum* | Prolate spheroidal | Loosely reticulated with small dots |
| *Plumeria alba* | Biconcave disc shaped | Smooth |
| *Portulaca grandiflora* | Prolate spheroidal | Spiny |
| *Sphagneticola trilobata* | Spherical | Spiny |
| *Tabernaemontana divaricata* | Biconcave disc shaped | Slightly rough |
| *Tagetes erecta* | Prolate spheroidal | Slightly rough |
| *Tamarindus indica* | Prolate spheroidal | Loosely reticulated |
| *Tecoma capensis* | Convex triangular | Densely reticulated |

**Fig. 2.**
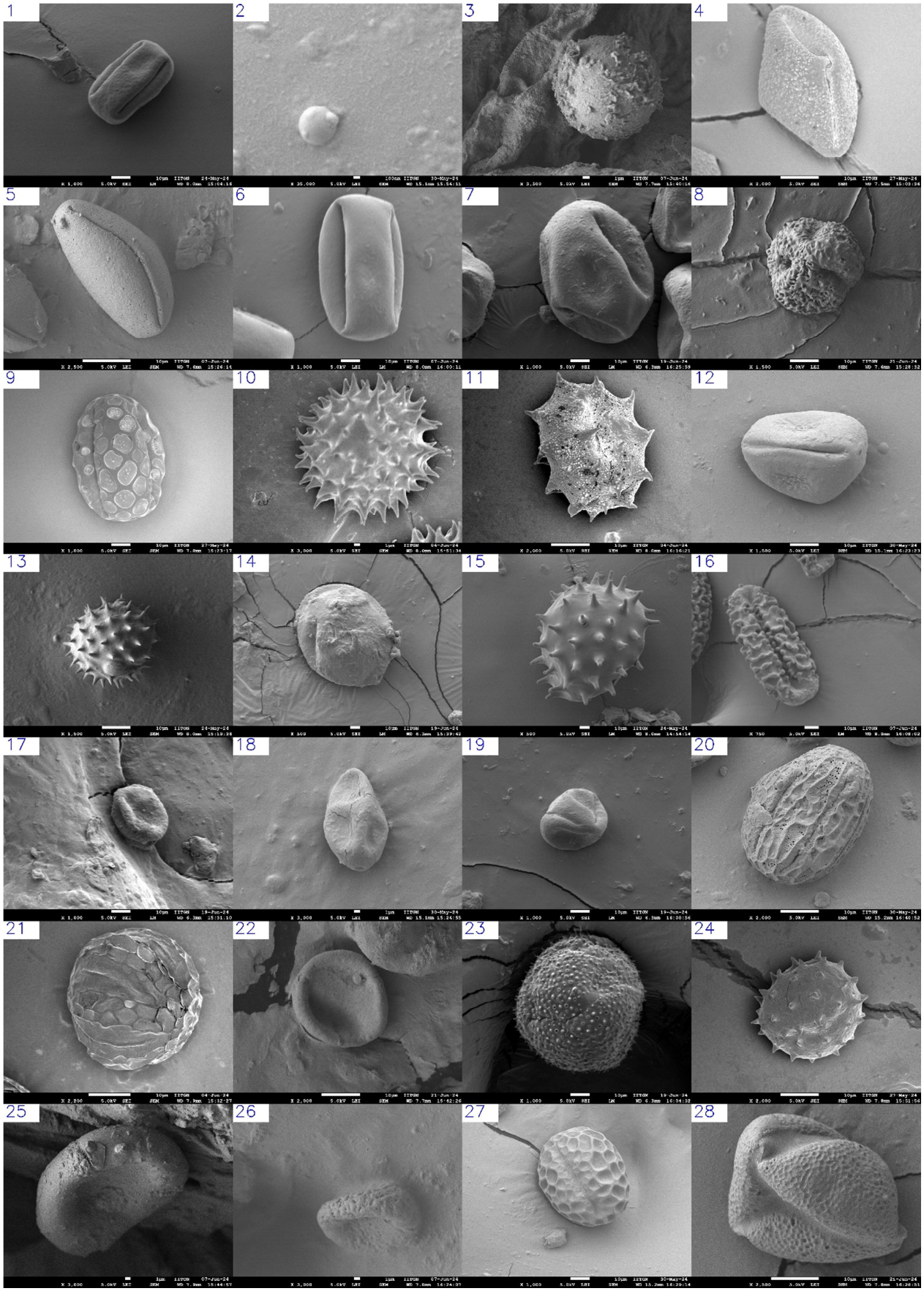
Pollen surface morphology of plant species used in this study. The pollen surface morphology of for plant speices such as 1) *Azadirachta indica*, 2) *Caesalpinia pulcherrima*, 3) *Canna indica*, 4) *Cassia fistula*, 5) *Cassia javanica*, 6) *Catharanthus pusillus*, 7) *Catharanthus roseus* (Linn.) Don, 8) *Cordia sebestena*, 9) *Delonix regia*, 10) *Eclipta prostrata*, 11) *Gaillardia pulchella*, 12) *Glandularia bipinnatifida*, 13) *Helianthus annuus*, 14) *Hibiscus arnottianus*, 15) *Hibiscus rosa-sinensis*, 16) *Hymenocallis littoralis*, 17) *Ixora coccinea*, 18) *Jasminum sambac*, 19) *Murraya paniculata*, 20) *Ocimum gratissimum*, 21) *Ocimum tenuiflorum*, 22) *Plumeria alba*, 23) *Portulaca grandiflora*, 24) *Sphagneticola trilobata*, 25) *Tabernaemontana divaricata*, 26) *Tagetes erecta*, 27) *Tamarindus indica*, 28) *Tecoma capensis* were determined using FE-SEM.

### IMPORTANT database for pollen SEM images

The web application developed for a systematic and structured display of pollen images and their microscopic features can be accessed through the IMPORTANT database (https://importantdb.pythonanywhere.com). This database provides details for the 28 flower species used in this study, including their traditional name, common name, kingdom, family, and description of the species. The SEM images for the pollen for the species can be searched through either the species name in the menu or the images for flowers in the gallery section. The user interface for the application is shown in **Fig 3**.

**Fig. 3.**
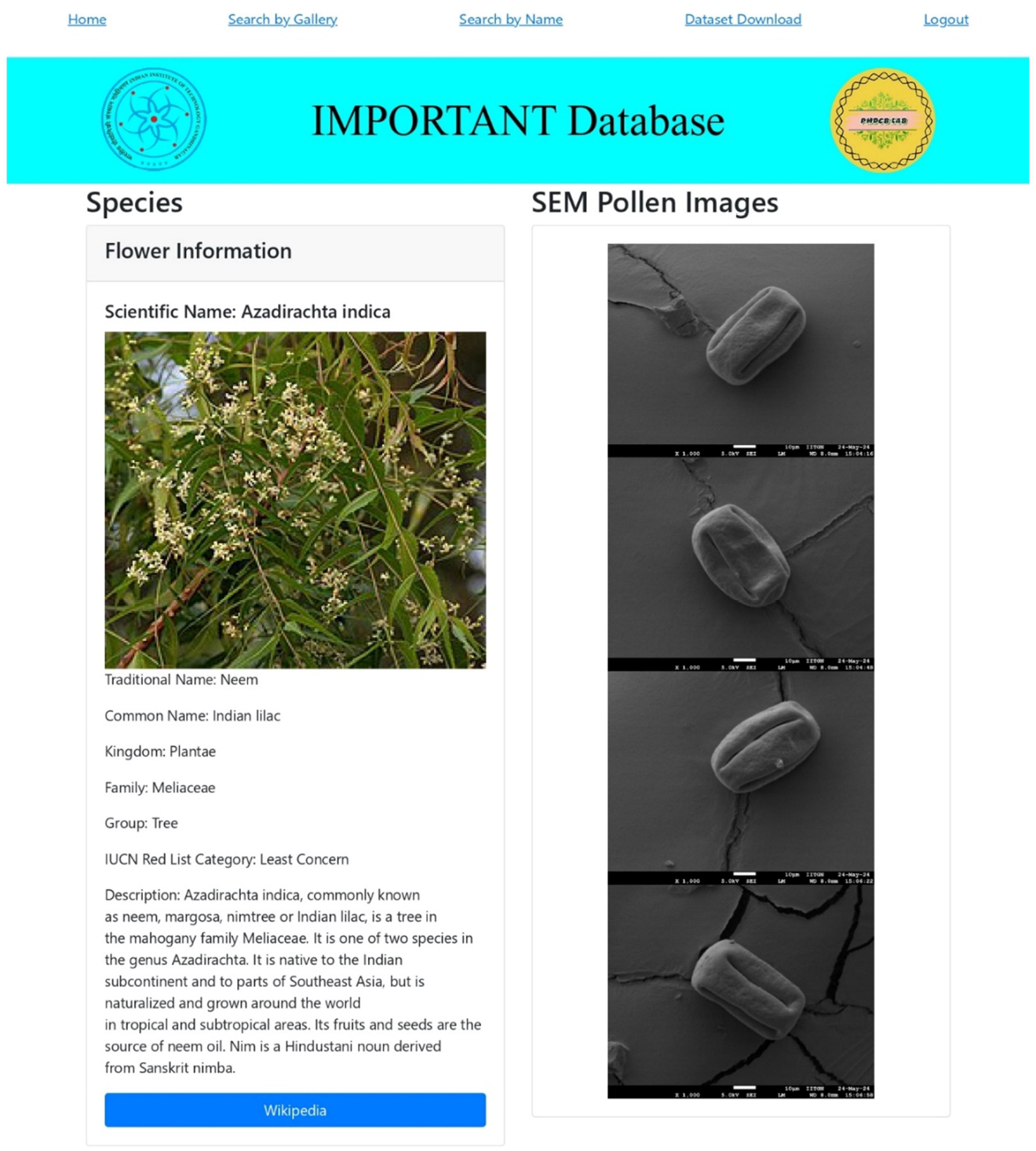
The user interface for IMPORTANT database. The IMPORTANT (Indian Medicinal Plants Pollen Image Database for Research, Training, Analysis, and Neural Networks Testing) database (https://importantdb.pythonanywhere.com) consists of the details of the pollen SEM images of 28 species used in this study. The pollen morphology for these plant species can be studied by searching the name or by clicking the image of the species.

### Segmentation and classification efficiently distinguish the pollen images for species

We have used YOLOv11n for the segmentation of pollen. The input of the images, along with their annotations, when masked with the actual image, resulted in the clear pollen images as displayed in **Fig. 4**. This step is essential as it reduces the amount of background noise present in the image and, therefore, enables us to perform a more accurate classification task using specialized architectures. YOLOv11excels at instance segmentation by accurately identifying and segmenting individual objects within images. It works with higher mean Average Precision (mAP) with fewer parameters and is adaptable across various environments for the instance segmentation task.YOLOv11n is built upon the advancements of its predecessors in the YOLO family and incorporates architectural enhancements, including improved backbone and neck architecture, facilitating feature extraction capabilities, and optimized training pipelines with enhanced processing speed. YOLO has been efficiently used in agriculture, including the detection of disease in tea and post-harvest physiological deterioration in cassava [30,31,32]. The box loss and the segmentation loss during the training phase decrease as we move ahead with more epochs, which results in an increase in precision and recall for both the box and the mask (**Fig. 5**). The details for the variation in multiple training results against the number of iterations during the training are detailed in **Fig. 5**. Since this was a single-class segmentation, the precision achieved for the box by segmentation model was 0.685, and the recall was 0.625. The mask precision achieved by the segmentation model was 0.689, while the mask recall was 0.629 (**Fig. 6**). The box corresponds to the blue colored bounding box, while the mask is the blue shaded region over the grayscale image of the pollen. The results showed that the bounding box is formed almost perfectly in most cases, while the mask is not perfect in some images (**Fig. 6**). This may be because of the inaccuracies of the model in generating the mask. The results of the segmentation model are provided in **Fig. 6**. The segmentation process proved to be highly efficient in isolating individual pollen grains from the complex background of SEM images. The automated segmentation method not only improved the speed of image analysis but also reduced human error associated with manual segmentation. This efficiency is particularly beneficial for large-scale studies involving extensive pollen sample collections.

**Fig. 4.**
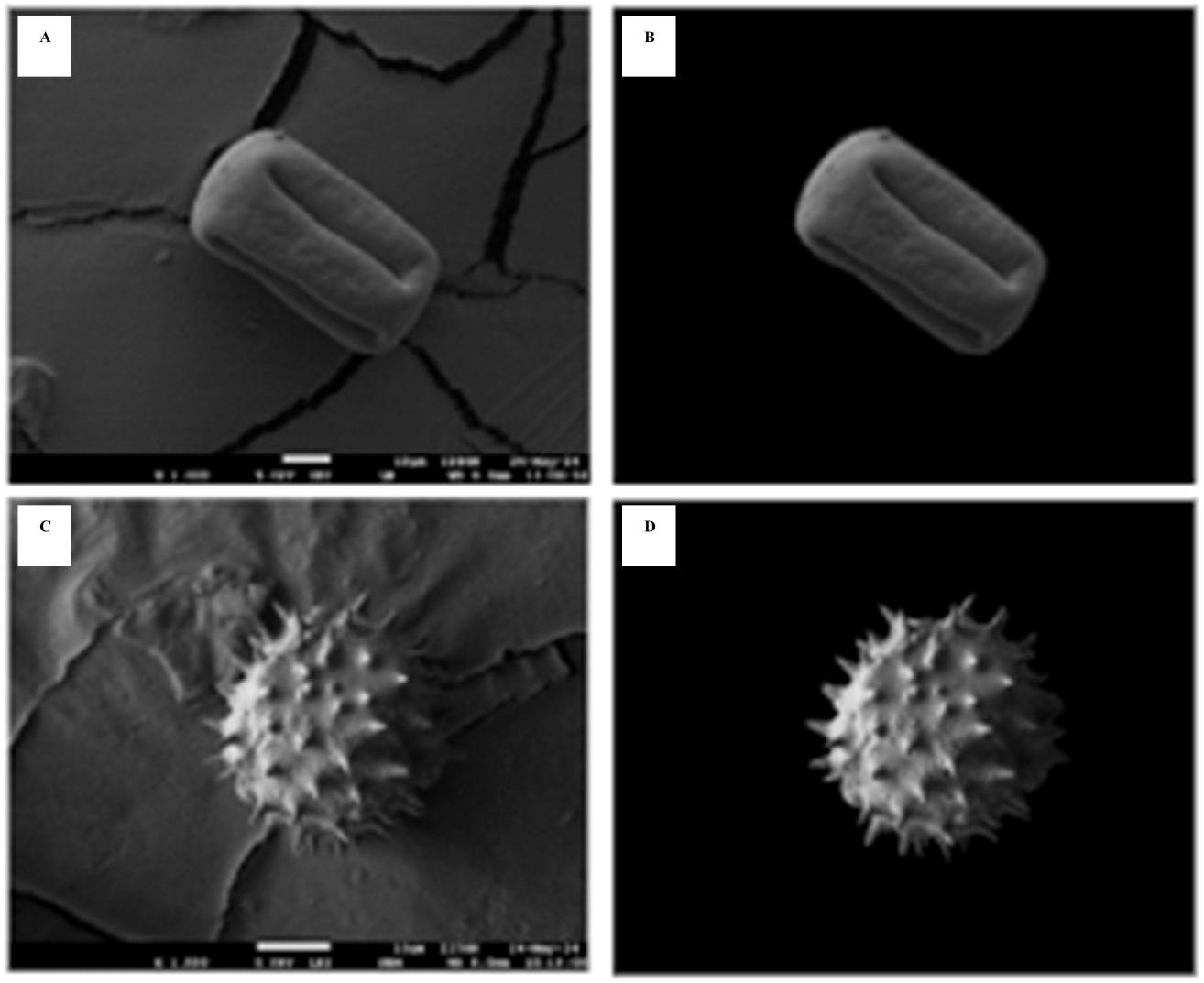
Segmented pollen images with annotated mask. The images A and C represent the pollen of *Azadirachta indica* and *Helianthus annuus* in the raw form with the unsegmented background. Images B and D are the pollen images segmented out from the background using manual annotation masks, which were used for training the segmentation models.

**Fig. 5.**
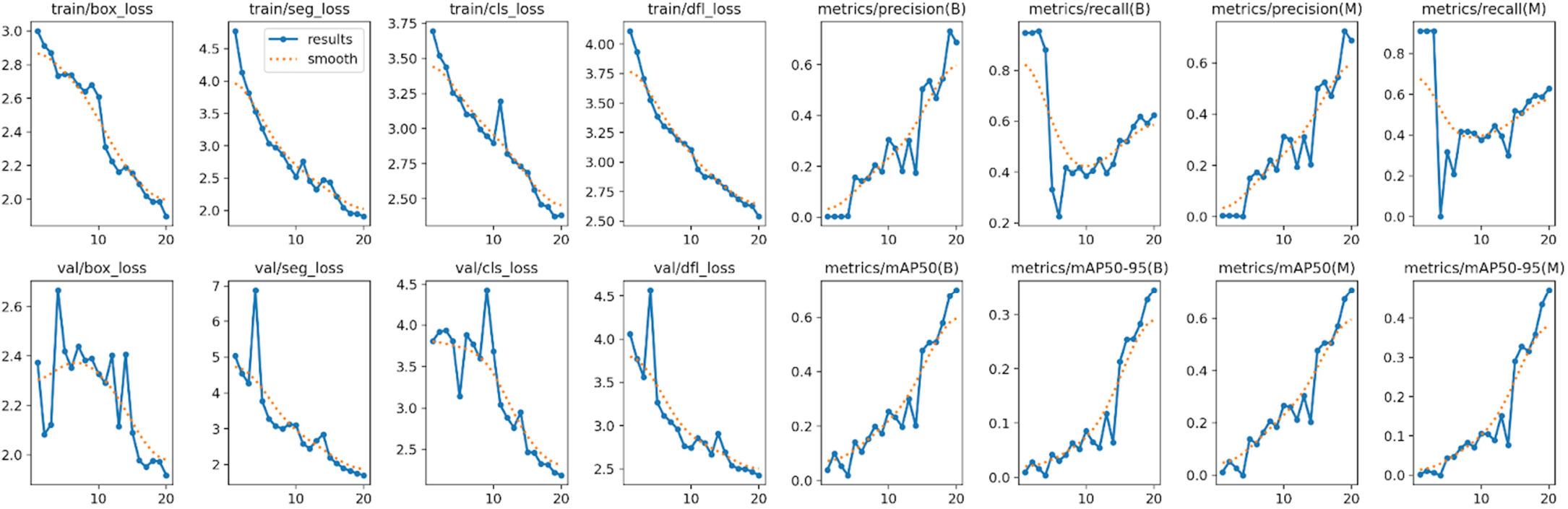
Training history plots for the YOLOv11n Segmentation model. The plots in the figure display the variation of multiple parameters against the number of epochs during the training phase for the YOLOv11n segmentation model.

**Fig. 6.**
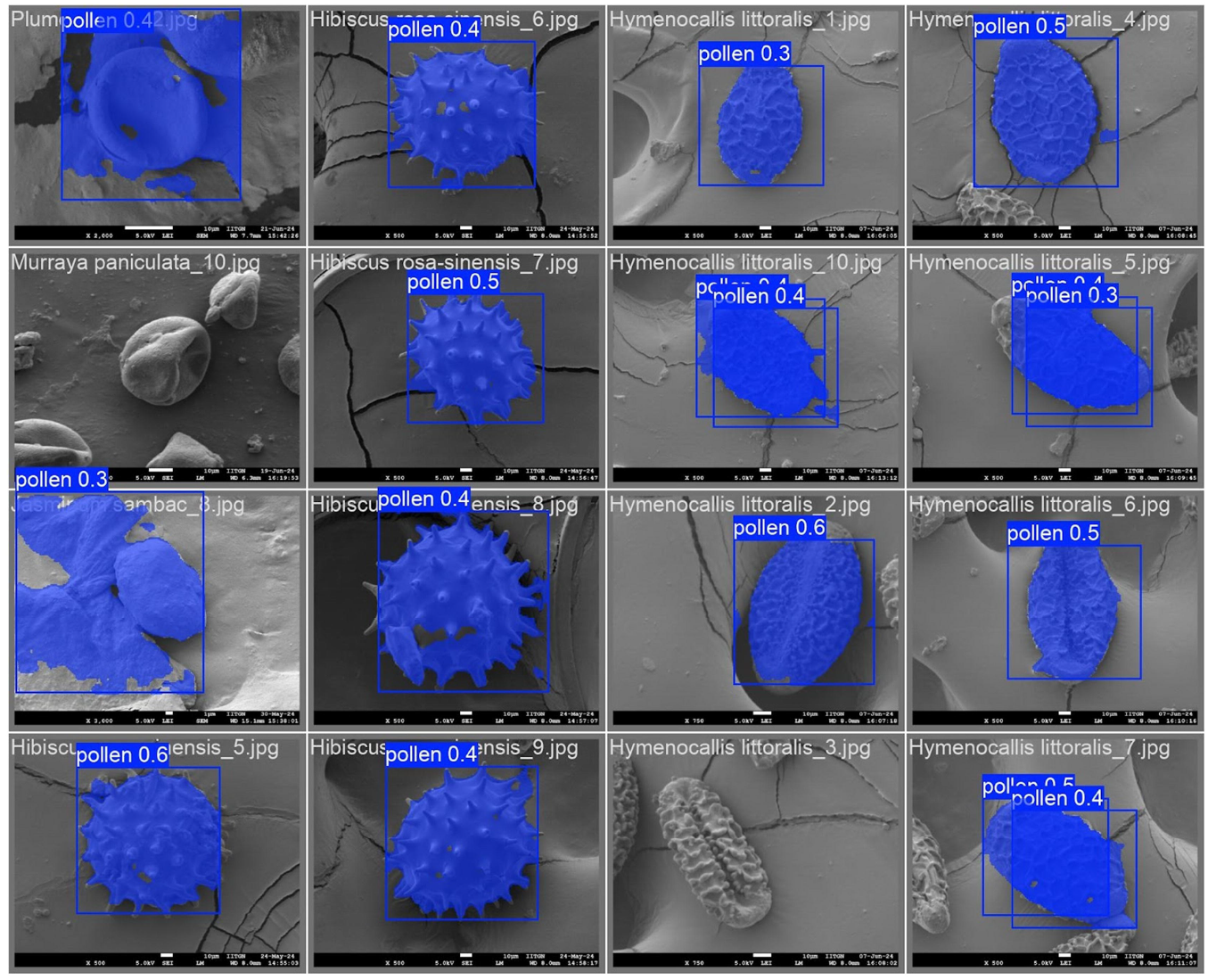
Mask and bounding box prediction by the Segmentation Model. Images display the bounding box as well as the segmentation mask on the test dataset predicted by the YOLOv11n model trained on the segmentation dataset.

In the classification of training models, most of the models perform quite well on the dataset, with the MobileNet model achieving the highest accuracy of 98% as well as the highest precision, recall and F1 score compared to other CNN-based models, suggesting the effectiveness of using CNN-based networks for the classification task on our dataset. MobileNet achieves a balance between latency, accuracy, and model size by using depth wise separable convolutions. This technique splits the convolution into two separate layers: a depth wise convolution that applies a single filter to each input channel and a pointwise convolution, efficiently combining the depth wise layer outputs. This approach is efficient over traditional convolutions as it reduces the computational complexity and parameter numbers, transforming it into a lightweight and efficient model.

In this study, the transformer model (ViT) had 100% accuracy in classification. ViT is one of the latest models that work on the principle of self-attention layers for image classification. The efficiency of this model can be determined from the fact that this model is pre-trained on the extensive ImageNet-21k dataset, which includes 14 million images and over 21,843 classes, which helps this model gain a robust understanding of diverse visual patterns. It can be efficiently used and fine-tuned for image classification, object detection, and segmentation. The ViT model is versatile and efficient in learning detailed images and uses approximately 30.1 million parameters and a 224×224 pixels processed image size, making it essential in various image-related applications. The detailed comparison between the performances of the models in terms of accuracy, macro average precision, macro average recall, and macro average F1 scores are provided in **Table 2**, and the classification report, comprising class-wise precision, recall, and F1 scores is provided in the **Supplementary Table. 4**.

**Table 2.** Comparison between the performances of various models on the dataset.

| Model | Accuracy | Macro Average Precision | Macro Average Recall | Macro Average F1 Score |
| --- | --- | --- | --- | --- |
| AlexNet | 0.71 | 0.69 | 0.68 | 0.65 |
| VGG 16 | 0.85 | 0.88 | 0.84 | 0.84 |
| ResNet 50 | 0.97 | 0.97 | 0.97 | 0.96 |
| MobileNet | 0.98 | 0.98 | 0.97 | 0.98 |
| ViT | 1.00 | 1.00 | 1.00 | 1.00 |

The results for the training history plots consisting of the training and validation accuracies against the number of epochs are given in **Fig. 7**. The CNN-based training history plot models (**Fig. 7a**), training history (accuracy) plot of the Vision Transformer-based Model (ViT) (**Fig. 7b**), and training history plot (loss) of the ViT Model (**Fig. 7c**) suggested that the accuracies during the training phase increased steadily for all the models, with ViT and ResNet50 saturating at higher accuracies at much earlier epochs compared to the other models, which gradually improved in the subsequent epochs. Though the ViT model saturated to a very high frequency in the early epochs, it continued to reduce the training loss over the subsequent epochs. The confusion matrix for the ViT model displays perfect classification since the non-diagonal values in the confusion matrix are exactly zero, while those of MobileNet and ResNet50 display slight inaccuracies (**Fig. 8** and **9**). Many of the classes were confused by the AlexNet model, making it the least accurate of the models under consideration for the classification task (**Fig. 8**). The confusion matrices for the test dataset consisting of 489 images are provided in **Fig. 8** and **9**.

**Fig. 7.**
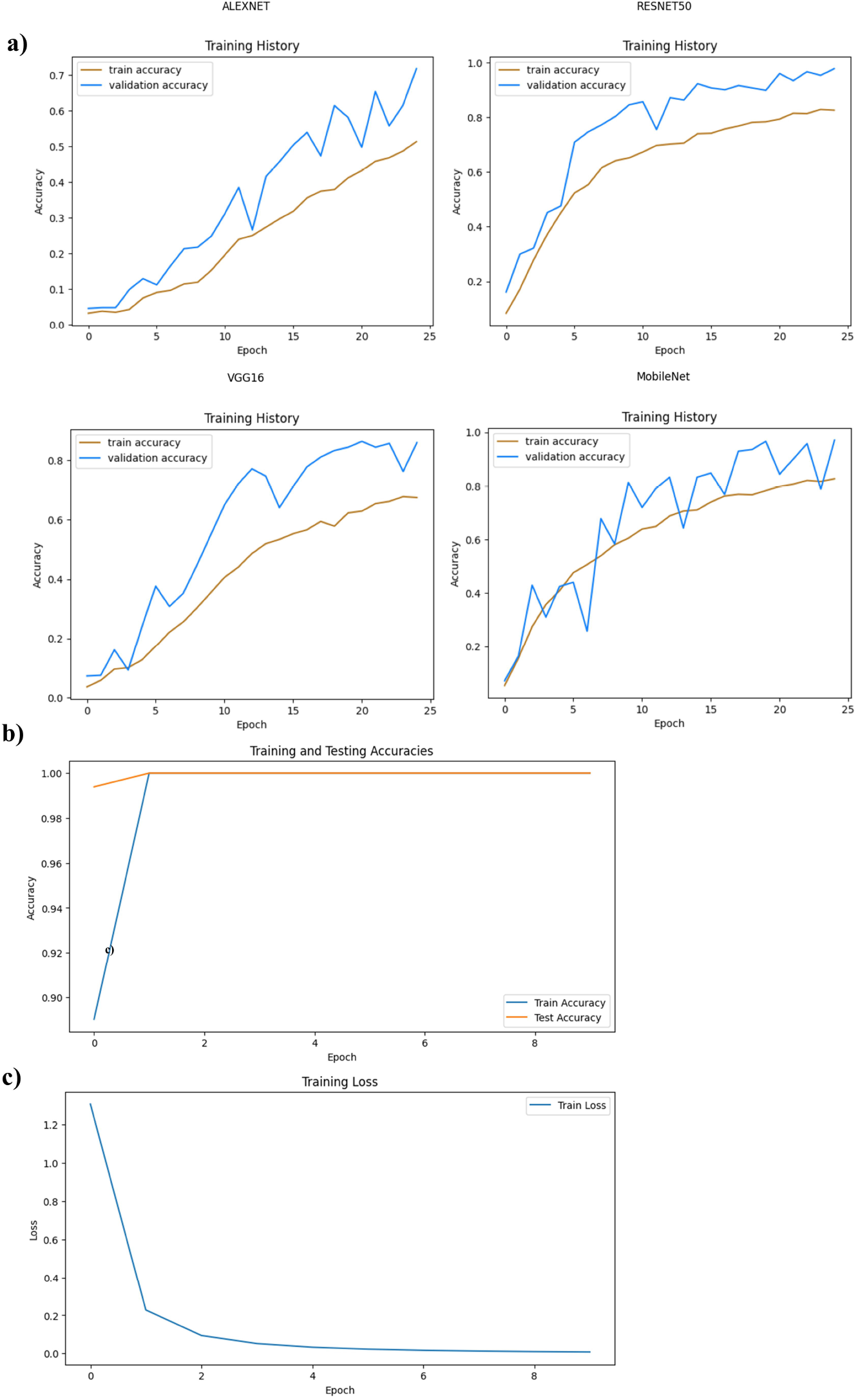
Training history plots showing training and validation accuracies versus the number of epochs. The training history plots for the model included a) CNN-based training history plot models, a) Training history (accuracy) plot of the Vision Transformer-based Model (ViT), and c) Training history plot (loss) of the ViT Model.

**Fig. 8.**
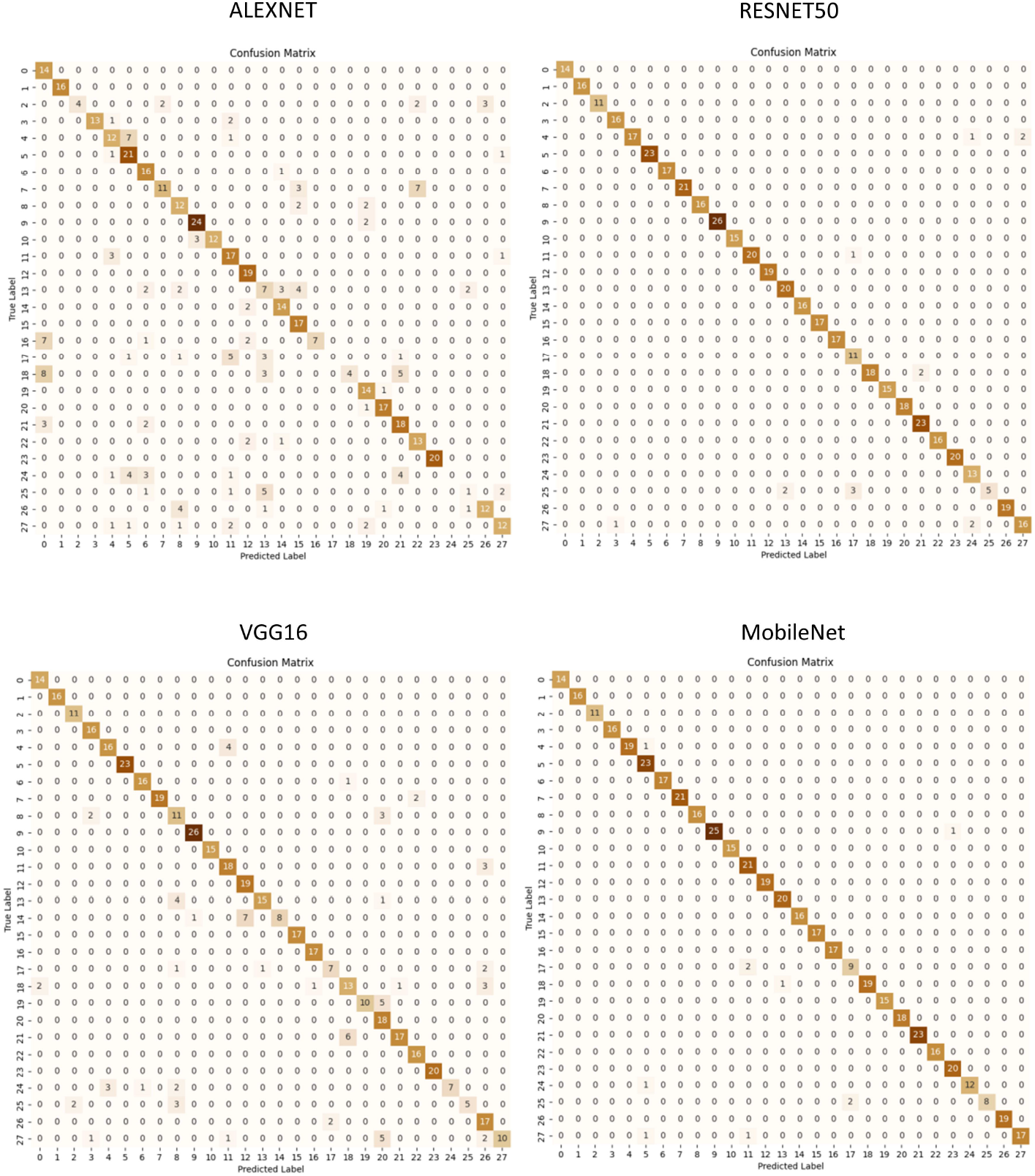
Confusion matrices of the CNN-based models. The plot displays the class-wise performance of the models (AlexNet, ResNet50, VGG16, and MobileNet) using the confusion matrix with the vertical axis representing the true label and the horizontal axis representing the predicted labels.

**Fig. 9.**
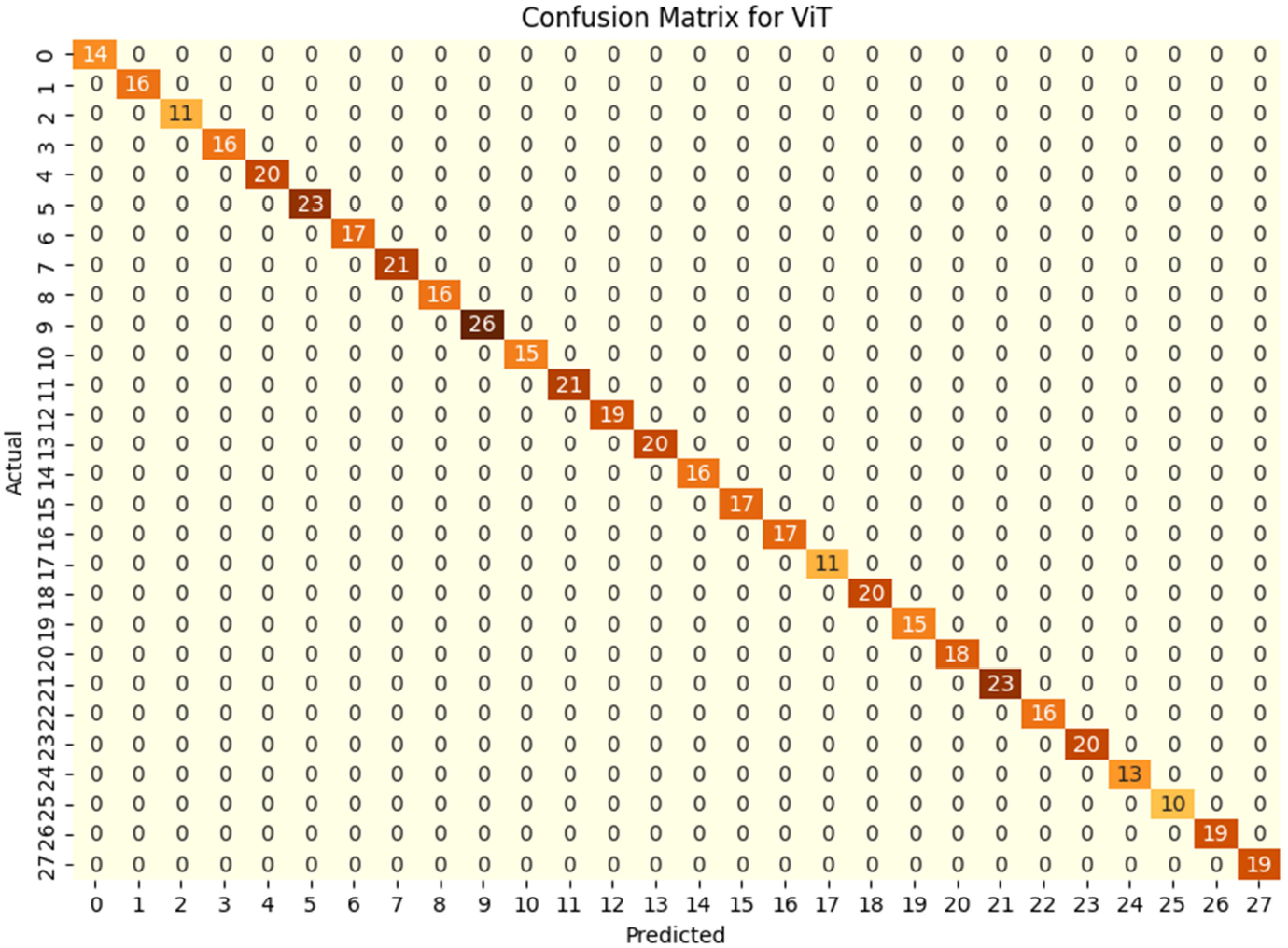
Confusion matrices of the Vision Transformer-based Model (ViT). The plot displays the class-wise performance of the Vision Transformer-based model (ViT) using the confusion matrix, with the vertical axis representing the true label and the horizontal axis representing the predicted labels.

In summary, the advancements in pollen image analysis using SEM and machine learning classification approach has various applications. The accuracy and precision of pollen identification are critical and have wide applications, including studies related to pollen allergy, historical climate changes, and archaeology. The dataset and methods developed in this study would offer a valuable tool for researchers providing a reliable and scalable approach to identification of pollen and medicinal plant species. However, our approach may have a few limitations, including capturing the details of morphological characters with high-quality SEM images, which may not be possible for some researchers due to limited access to research facilities. In addition, there is always a possibility of error in the misclassification by classification model, mostly in the case of morphologically similar intra-related species. The development of the segmentation models and algorithms for the pollen images can effectively address these issues.

## Conclusion

The integration of SEM imaging with advanced classification and segmentation techniques represents a significant advancement in pollen research. This research contributes to the pollen SEM image dataset consisting of 269 images and their annotations in 28 classes for the segmentation task and 5842 images in 28 classes for the classification task, along with the development of an extensive database with a user-friendly interface. The main highlight of this dataset is its focus on the pollen of common medicinal plant species found in India, which have not been explored previously. The future scope of this project is to expand the dataset to cover more species and integrate this at a software level with the SEM to automate the task of segmentation and classification of the pollen directly from the SEM images, leading to the development of efficient segmentation and classification models that will be developed in the future. This research will not only benefit scientific research but also have practical applications in various fields such as agriculture and archaeological science. Overall, our work will be instrumental in future advancements in pollen research, especially in Indian medicinal plants.

## Supplementary information

Three Supplementary Tables 1 to 4 are provided with the manuscript.

## Declarations

### Funding

This work was supported by a DBT Ramalingaswami Re-entry fellowship grant and a start-up grant from the Indian Institute of Technology Gandhinagar to S.S. J.S.K. was supported by an SRIP fellowship from IITGN. N.D.G. is supported by a post-doctoral fellowship from IITGN.

### Conflict of interest

The authors declare no competing interests

### Ethics approval and consent to participate

Not applicable

### Consent for publication

Not applicable

### Data availability

IMPORTANT Database (https://importantdb.pythonanywhere.com)

### Materials availability

Not applicable

### Author contribution

S.S. and J.S.K. conceived and designed the research. J.S.K. performed all of the experiments, and N.D.G. assisted in SEM. S.R. provided technical suggestions for the experiments. S.S. and S.R. supervised the experiments. J.S.K. wrote the first draft of the manuscript. N.D.G., S.S., and S.R. proofread and edited the manuscript. All authors read, discussed and approved the manuscript.

## Acknowledgments

We thank the Indian Institute of Technology Gandhinagar for the internship opportunity at JSK and the fellowship to NG. We also thank the Central Instrumentation Facility (CIF) at IITGN for the access to SEM.

## Notes

### Competing Interest Statement

The authors have declared no competing interest.

