## Supplementary Tables for "IMPORTANT: Advanced Pollen Classification of Indian Medicinal Plants through SEM and Computer Vision"

**Supplementary Table 3**. Distribution of classification dataset across species

| Sr. No. | Plant species | Number of images used for classification |
| --- | --- | --- |
| 1 | *Azadirachta indica* | 176 |
| 2 | *Caesalpinia pulcherrima* | 198 |
| 3 | *Canna indica* | 132 |
| 4 | *Cassia fistula* | 198 |
| 5 | *Cassia javanica* | 220 |
| 6 | *Catharanthus pusillus* | 220 |
| 7 | *Catharanthus roseus* (Linn.) Don | 220 |
| 8 | *Cordia sebestena* | 220 |
| 9 | *Delonix regia* | 198 |
| 10 | *Eclipta prostrata* | 220 |
| 11 | *Gaillardia pulchella* | 220 |
| 12 | *Glandularia bipiinnatifida* | 220 |
| 13 | *Helianthus annuus* | 220 |
| 14 | *Hibiscus arnottianus* | 242 |
| 15 | *Hibiscus rosa-sinensis* | 220 |
| 16 | *Hymenocallis littoralis* | 220 |
| 17 | *Ixora coccinea* | 176 |
| 18 | *Jasminum sambac* | 176 |
| 19 | *Murraya paniculata* | 220 |
| 20 | *Ocimum gratissum* | 242 |
| 21 | *Ocimum tenuiflorum* | 220 |
| 22 | *Plumeria alba* | 220 |
| 23 | *Portulaca grandiflora* | 220 |
| 24 | *Sphagneticola trilobata* | 220 |
| 25 | *Tabernaemontana divaricata* | 154 |
| 26 | *Tagetes erecta* | 88 |
| 27 | *Tamarindus indica* | 242 |
| 28 | *Tecoma capensis* | 220 |
|  | Total | 5842 |

Note: In the table, P denotes precision, R is recall, and F1 is for F1 score.
