## Supplementary Tables for "IMPORTANT: Advanced Pollen Classification of Indian Medicinal Plants through SEM and Computer Vision"

**Supplementary Table 4**. Classification reports for various architectures for the models

| Plant species | AlexNet | | | MobileNet | | | ResNet 50 | | | VGG 16 | | |
| --- | --- | --- | --- | --- | --- | --- | --- | --- | --- | --- | --- | --- |
|  | P | R | F1 | P | R | F1 | P | R | F1 | P | R | F1 |
| *Azadirachta indica* | 0.44 | 1 | 0.61 | 1 | 1 | 1 | 1 | 1 | 1 | 0.88 | 1 | 0.93 |
| *Caesalpinia pulcherrima* | 1 | 1 | 1 | 1 | 1 | 1 | 1 | 1 | 1 | 1 | 1 | 1 |
| *Canna indica* | 1 | 0.36 | 0.53 | 1 | 1 | 1 | 1 | 1 | 1 | 0.85 | 1 | 0.92 |
| *Cassia fistula* | 1 | 0.81 | 0.9 | 1 | 1 | 1 | 0.94 | 1 | 0.97 | 0.84 | 1 | 0.91 |
| *Cassia javanica* | 0.63 | 0.6 | 0.62 | 1 | 0.95 | 0.97 | 1 | 0.85 | 0.92 | 0.84 | 0.8 | 0.82 |
| *Catharanthus pusillus* | 0.62 | 0.91 | 0.74 | 0.88 | 1 | 0.94 | 1 | 1 | 1 | 1 | 1 | 1 |
| *Catharanthus roseus (Linn.) Don* | 0.64 | 0.94 | 0.76 | 1 | 1 | 1 | 1 | 1 | 1 | 0.94 | 0.94 | 0.94 |
| *Cordia sebestena* | 0.85 | 0.52 | 0.65 | 1 | 1 | 1 | 1 | 1 | 1 | 1 | 0.9 | 0.95 |
| *Delonix regia* | 0.6 | 0.75 | 0.67 | 1 | 1 | 1 | 1 | 1 | 1 | 0.52 | 0.69 | 0.59 |
| *Eclipta prostrata* | 0.89 | 0.92 | 0.91 | 1 | 0.96 | 0.98 | 1 | 1 | 1 | 0.96 | 1 | 0.98 |
| *Gaillardia pulchella* | 1 | 0.8 | 0.89 | 1 | 1 | 1 | 1 | 1 | 1 | 1 | 1 | 1 |
| *Glandularia bipiinnatifida* | 0.59 | 0.81 | 0.68 | 0.88 | 1 | 0.93 | 1 | 0.95 | 0.98 | 0.78 | 0.86 | 0.82 |
| *Helianthus annuus* | 0.76 | 1 | 0.86 | 1 | 1 | 1 | 1 | 1 | 1 | 0.73 | 1 | 0.84 |
| *Hibiscus arnottianus* | 0.37 | 0.35 | 0.36 | 0.95 | 1 | 0.98 | 0.91 | 1 | 0.95 | 0.94 | 0.75 | 0.83 |
| *Hibiscus rosa-sinensis* | 0.74 | 0.88 | 0.8 | 1 | 1 | 1 | 1 | 1 | 1 | 1 | 0.5 | 0.67 |
| *Hymenocallis littoralis* | 0.65 | 1 | 0.79 | 1 | 1 | 1 | 1 | 1 | 1 | 1 | 1 | 1 |
| *Ixora coccinea* | 1 | 0.41 | 0.58 | 1 | 1 | 1 | 1 | 1 | 1 | 0.94 | 1 | 0.97 |
| *Jasminum sambac* | 0 | 0 | 0 | 0.82 | 0.82 | 0.82 | 0.73 | 1 | 0.85 | 0.78 | 0.64 | 0.7 |
| *Murraya paniculata* | 1 | 0.2 | 0.33 | 1 | 0.95 | 0.97 | 1 | 0.9 | 0.95 | 0.65 | 0.65 | 0.65 |
| *Ocimum gratissum* | 0.89 | 0.94 | 0.92 | 1 | 1 | 1 | 1 | 1 | 1 | 0.56 | 1 | 0.72 |
| *Ocimum tenuiflorum* | 0.67 | 0.93 | 0.78 | 1 | 1 | 1 | 1 | 1 | 1 | 1 | 0.67 | 0.8 |
| *Plumeria alba* | 0.64 | 0.78 | 0.71 | 1 | 1 | 1 | 0.92 | 1 | 0.96 | 0.94 | 0.74 | 0.83 |
| *Portulaca grandiflora* | 0.59 | 0.81 | 0.68 | 1 | 1 | 1 | 1 | 1 | 1 | 0.89 | 1 | 0.94 |
| *Sphagneticola trilobata* | 1 | 1 | 1 | 0.95 | 1 | 0.98 | 1 | 1 | 1 | 1 | 1 | 1 |
| *Tabernaemontana divaricata* | 0 | 0 | 0 | 1 | 0.92 | 0.96 | 0.81 | 1 | 0.9 | 1 | 0.54 | 0.7 |
| *Tagetes erecta* | 0.25 | 0.1 | 0.14 | 1 | 0.8 | 0.89 | 1 | 0.5 | 0.67 | 1 | 0.5 | 0.67 |
| *Tamarindus indica* | 0.8 | 0.63 | 0.71 | 1 | 1 | 1 | 1 | 1 | 1 | 0.63 | 0.89 | 0.74 |
| *Tecoma capensis* | 0.75 | 0.63 | 0.69 | 1 | 0.89 | 0.94 | 0.89 | 0.84 | 0.86 | 1 | 0.53 | 0.69 |

Note: In the table, P denotes precision, R is recall, and F1 is for F1 score.
